# Impulse model-based differential expression analysis of time course sequencing data

**DOI:** 10.1101/113548

**Authors:** David S. Fischer, Fabian J. Theis, Nir Yosef

**Affiliations:** Institute of Computational Biology, Helmholtz Centre Munich; TUM School of Life Sciences Weihenstephan, Technical University of Munich; Department of Mathematics, Technical University of Munich; Department of Electrical Engineering &Copmuter Science and Center for Computational Biology, University of California, Berkeley CA; Ragon Institute of MGH, MIT& Harvard, Cambridge MA

## Abstract

The global gene expression trajectories of cellular systems in response to developmental or environmental stimuli often follow the prototypic single-pulse or state-transition patterns which can be modeled with the impulse model. Here we combine the continuous impulse expression model with a sequencing data noise model in ImpulseDE2, a differential expression algorithm for time course sequencing experiments such as RNA-seq, ATAC-seq and ChIP-seq. We show that ImpulseDE2 outperforms currently used differential expression algorithms on data sets with sufficiently many sampled time points. ImpulseDE2 is capable of differentiating between transiently and monotonously changing expression trajectories. This classification separates genes which are responsible for the initial and final cell state phenotypes from genes which drive or are driven by the cell state transition and identifies down-regulation of oxidative-phosphorylation as a molecular signature which can drive human embryonic stem cell differentiation.

---

Time course sequencing experiments such as RNA-seq, ChIP-seq and ATAC-seq yield a description of the development of a cellular system over time. Such a dynamic description can be used to analyze the timing of cellular programs and can uncover transitional responses which are not observed if only initial and terminal cell states are compared. These dynamic properties give insights into the regulatory molecular circuits that drive the developmental process. Differential expression analysis is frequently used to reduces time course (longitudinal) data sets to genes with varying expression profiles across conditions to ease downstream analytic tasks.

Differential expression analysis algorithms for time course data sets can be divided into methods which model dependence between time points and those which treat time points independently. Independence of expression levels between time points can be modeled with generalized linear models based on time as a categorial (and not continuous) predictor of the expression level (which can be done with DESeq [1], DESeq2 [2], edgeR [3] and limma [4] for example). Constraining the sequence of inferred expression levels to a continuous function of time captures dependence of expression levels between time points. Examples for such functions previously used for differential expression analysis are natural cubic splines (edge [5]), polynomial models (maSigPro [6]), gaussian processes (DyNB [7]) and the impulse model (ImpulseDE [8]).

Categorial time models suffer from a relative loss of statistical testing power relative to expression models with a fixed number of parameters if many time points are observed. Moreover, categorial time models are prone to fit multi-modal low amplitude signals which may not be of interest to the analyst (Fig. 1D). Lastly, categorial time models are difficult to use if expression trajectories are compared between conditions which were sampled at different time points.

**Figure 1:**
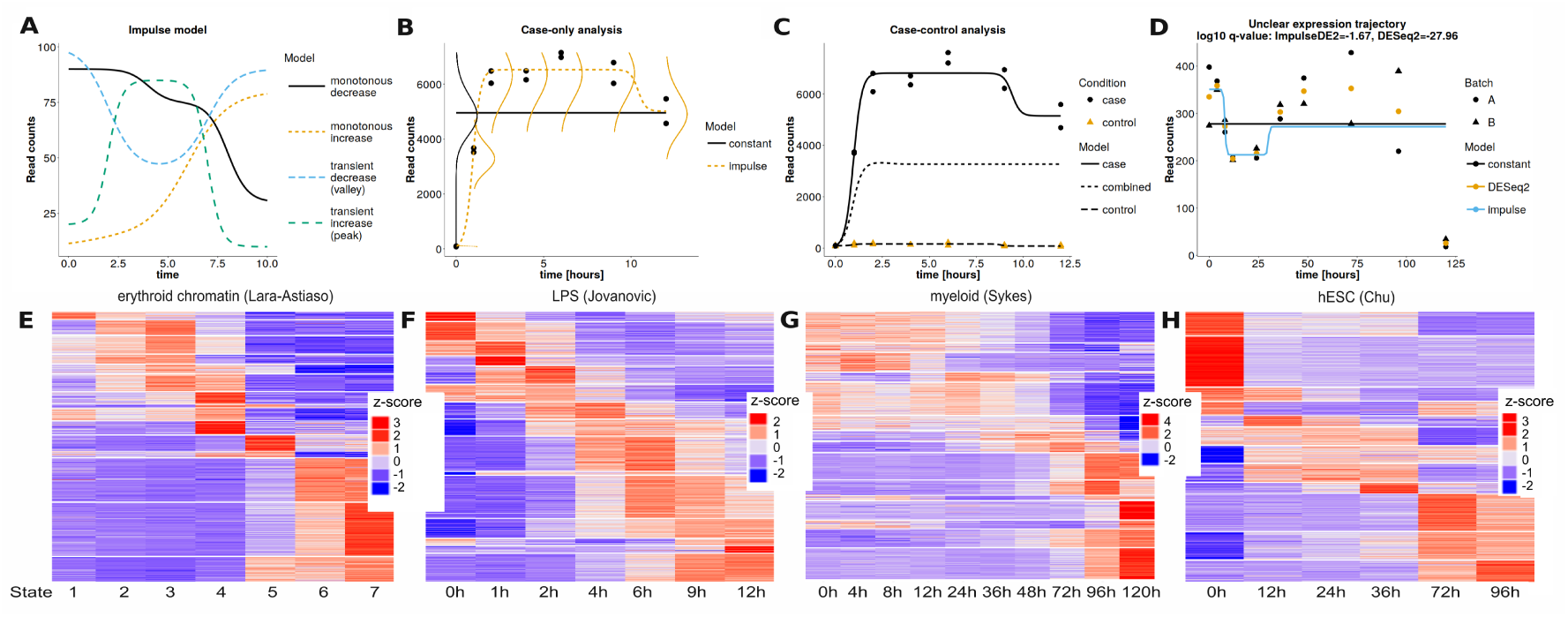
The impulse model is descriptive of global transcriptome and chromatin dynamics during the cellular response to stimuli. **A** The four classes of expression trajectories that can be modeled with the impulse model. **B** Case-only analysis: Shown are an impulse fit (alternative model) and a constant fit (null model) with vertically superimposed inferred negative binomial likelihood functions. The likelihood functions describe the likelihood of observing a read count given the mean expression level modeled by the impulse or constant and a dispersion factor. The likelihood functions are scaled and shifted so that the density is zero at the time coordinate of the time point of sampling. Low densities are not shown. **C** Case-control analysis: Shown are a separate case and control impulse fit (alternative model) and a single impulse fit to all samples (’’combined”, null model). **D** A potentially multi-modal expression trajectory from the myeloid (Sykes) data set. Shown are constant and impulse model fit by ImpulseDE2 (continuous models) and the fit of DESeq2 to time as a categorial variable: One value per observed time point. All fits are fits to batch A. Fits to batch B are the fits of batch A scaled by a single constant for this transcript. **E** Chromatin immunoprecipitation (ChIP) of the H3K4me1 histone mark in the erythroid lineage in hematopoesis (”erythroid chromatin (Lara-Astiaso)” data set [10]). The x-cordinates are seven cell states in developmental order (’developmental times”) within the erythroid lineage. We assigned uniformly spaced integers as times to the cell states. Rows are iChIP peaks called with MACS2 [16] and the signal is the number of reads overlapping a peak. Peaks were called on the union of reads of all samples of one cell type and the union of all peak sets of the cell types in the erythroid lineage was used as a reference based on which overlaps. **F** RNA-seq of dendritic cell activation through LPS (”LPS (Jovanovic)” data set [11]).**G** RNA- seq of myeloid differentiation (”myeloid (Sykes)” data set [13]).**H** RNA-seq of differentiation of human embryonal stem cells to definite endoderm (”hESC (Chu)” data set [14]). Refer to the Supplementary Methods for further details on the heatmaps. Heatmap of two further data sets (”estrogen (Baran-Gale)” [12] and ‘’Plasmodium (Broadbent)” data set [15]) are in Fig. SI1.

Continuous expression models of time can in principle overcome these shortcomings of categorial time models. However, these algorithms suffer from at least one of two pitfalls: Firstly, the underlying continuous models are limited in their ability to fit the temporal profiles in molecular data sets. For instance, DyNB uses a gaussian process, which may either over-fit multi-modal signals (similar to categorial time models) or miss drastic expression level transitions which we often observe in sequencing data (Fig. 1E-H). Secondly, the statistical model used are not adequate for capturing the underlying process of read generation. As a result, categorial models of time are still used widely for the analysis of time course data, primarily because of their convergence properties, advanced noise models developed specifically for sequencing count data, such as in DESeq2 [2] (negative binomial noise and dispersion trend smoothing) and their robustness to signal shape.

Here, we present ImpulseDE2, a differential expression algorithm for longitudinal sequencing experiments. ImpulseDE2 models the gene-wise expression trajectories over time with a descriptive single-pulse (impulse) function. The impulse function represents deviation from a steady initial state to a transition state and return to a new steady state (Fig. 1A). As we demonstrate below, the impulse function fits a large range of cellular responses to environmental or developmental cues 1E-H) [9]. We have previously utilized the impulse pattern as a part of the differential expression framework ImpulseDE. ImpulseDE2 combines the descriptiveness of the impulse model with a noise model tailored to sequencing data, which builds on and extends the DESeq2 model.

Here, we show that ImpulseDE2 over-comes the short-comings of categorial time models without acquiring the pitfalls of continuous expression models in time (convergence problems and low descriptiveness). Moreover, we show that the adapted DESeq2 noise model improves performance of ImpulseDE2 versus differential expression algorithms based on other noise models and versus DESeq2 itself.

Secondly, we show that the impulse model can be used to annotate differentially expressed genes with a rough description of their role in the cellular response. We introduce a hypothesis testing scheme that compares a constant, with a sigmoid and an impulse fit of each gene. This comparison can be used to identify transiently or permanently activated or deactivated genes. These classes of transient and permanent changes directly relate to the biological process of cell activation or differentiation: Over the course of the response of a cell to a stimulus, the cell moves from one state in the transcriptome space to another state. ImpulseDE2 can distinguish genes responsible for the differences between the states (permanent changes) and genes which change transiently during the transition between the cell states.

ImpulseDE2 is implemented as a R package and available through GitHub at https://github.com/YosefLab/ImpulseDE2.

## 1 Results

ImpulseDE2 is a differential expression algorithm which combines a negative binomial noise model (Fig. 1B) with a uni-modal parametric model for the expression trajectory across time (impulse model, Fig. 1A). ImpulseDE2 fits expression trajectories numerically based on DESeq2 [2] dispersion estimates set as hyperparameters and evaluates differential expression based on a log-likelihood ratio test. ImpulseDE2 combines the strengths of a descriptive expression models continuous in time with the noise modeling adapted from DESeq2. Algorithmic novelties include a non-standard batch model for case-control comparison, automated DESeq2 dispersion outlier handling and maximum likelihood-based fitting of the impulse model based on a negative binomial noise model.

ImpulseDE2 has two modes of operation: Single-condition differential expression analysis (”case-only”) and two-condition differential expression analysis (”case-control”). Case-only differential expression analysis identifies genes which have non-constant expression trajectories over time from samples of a single condition (Fig. 1B). Case-control differential expression analysis identifies genes which have different expression trajectories over time between two conditions (such as with and without a stimulus or treatment at time point zero) (Fig. 1C).

We compared ImpulseDE2 with the following reference methods: DESeq2, based on a categorial time expression model and with a negative binomial noise model, edge, based on natural cubic splines as expression models with a model-free error model and ImpulseDE, based on the impulse model as the expression model with a model-free error model.

### 1.1 The impulse model is descriptive of global transcriptome and chromatin dynamics during the cellular response to stimuli

We selected differentially expressed genes without constraints on the expression trajectory with DESeq2 and clustered the expression profiles of individual genes over time (Fig. 1E-H, SI1). All shown clusters can be modeled with the impulse model as “peaks” or ‘’valleys” and a few exceptional clusters with weak bi-modal behavior can be approximated with a uni-modal model. The presented data sets cover histone mark dynamics during development [10] (Fig. 1E), expression profiles of cell cultures in responseto environmental stimuli [11] [12] (Fig. 1F, SI1A) and expression profiles of cell cultures in response to developmental stimuli [13] [14] [15] (Fig. 1G,H, SI1B).

### 1.2 ImpulseDE2 has advantages over DESeq2 and edge on simulated data

We simulate gene-wise expression trajectories based on constant, linear, sigmoid and impulse models parameterized randomly for each gene to give trajectories within a target count range. The simulated data sets are designed to contain roughly equally many differentially and non-differentially expressed genes out of 6000 genes simulated for each data set: In case-only differential expression analysis, non-differentially expressed genes are simulated with constant trajectories, differentially expressed genes with linear, sigmoid or impulse trajectories. In case-control differential expression analysis, non differentially expressed genes are simulated with the same functional form and the same parameterizations for each condition and gene, differently expressed genes with the same functional form but different parameterizations for each condition and gene. On top of the simulated expression trajectories, we simulate size factors, batch effects and negative binomial noise as assumed in the DESeq2 and ImpulseDE2 noise model. ImpulseDE and edge have a model free noise model but both assume independent and identically distributed noise. Therefore, we log-transformed size-factor normalized count data as input for ImpulseDE and edge. 6000 genes were simulated for each data set. A detailed description of the simulation can be found in the Supplementary Methods.

We measure method performance with the area-under the curve (AUC) of the receiver-operator characteristic (ROC) curve for which each differential expression method is viewed as a classifier that associates a label (differentially expressed or not differentially expressed) with each gene. The classifier boundary can be moved by shifting the threshold p-value for calling differential expression. We also attach precision-recall statistics in the supplement (Fig. SI3). We analyze p-values as oppose to false-discovery rate (FDR) corrected q-values as different FDR corrections are implemented for the methods described.

As expected, ImpulseDE2 and edge have higher statistical power and therefore higher AUC of the ROC curve than the categorial time model DESeq2 if many time points are sampled in both the case-only and the case-control scenario (Fig. 2A,B). ImpulseDE2 and edge have a consistently lower false discovery rate than DESeq2 at different levels of noise on randomly generated expression trajectories (Fig. 2C). ImpulseDE2 and edge can perform case-control differential expression analysis similarly well if both conditions are sampled at the time points or at different time points (asynchronous sampling)(Fig. 2D). Non-synchronous sampling may arise with patient data and cannot be easily handled with DESeq2. ImpulseDE2 does not suffer from numerical problems in batch effect correction: ImpulseDE2 and DESeq2 perform similarly well on data with batch effects (Fig. 2E), which is expected because their batch models do not differ in the simulated scenarios. The model-free error model of edge is more susceptible to identify constant expression trajectories as differentially expressed at low expression levels (below 20 counts) than

**Figure 2:**
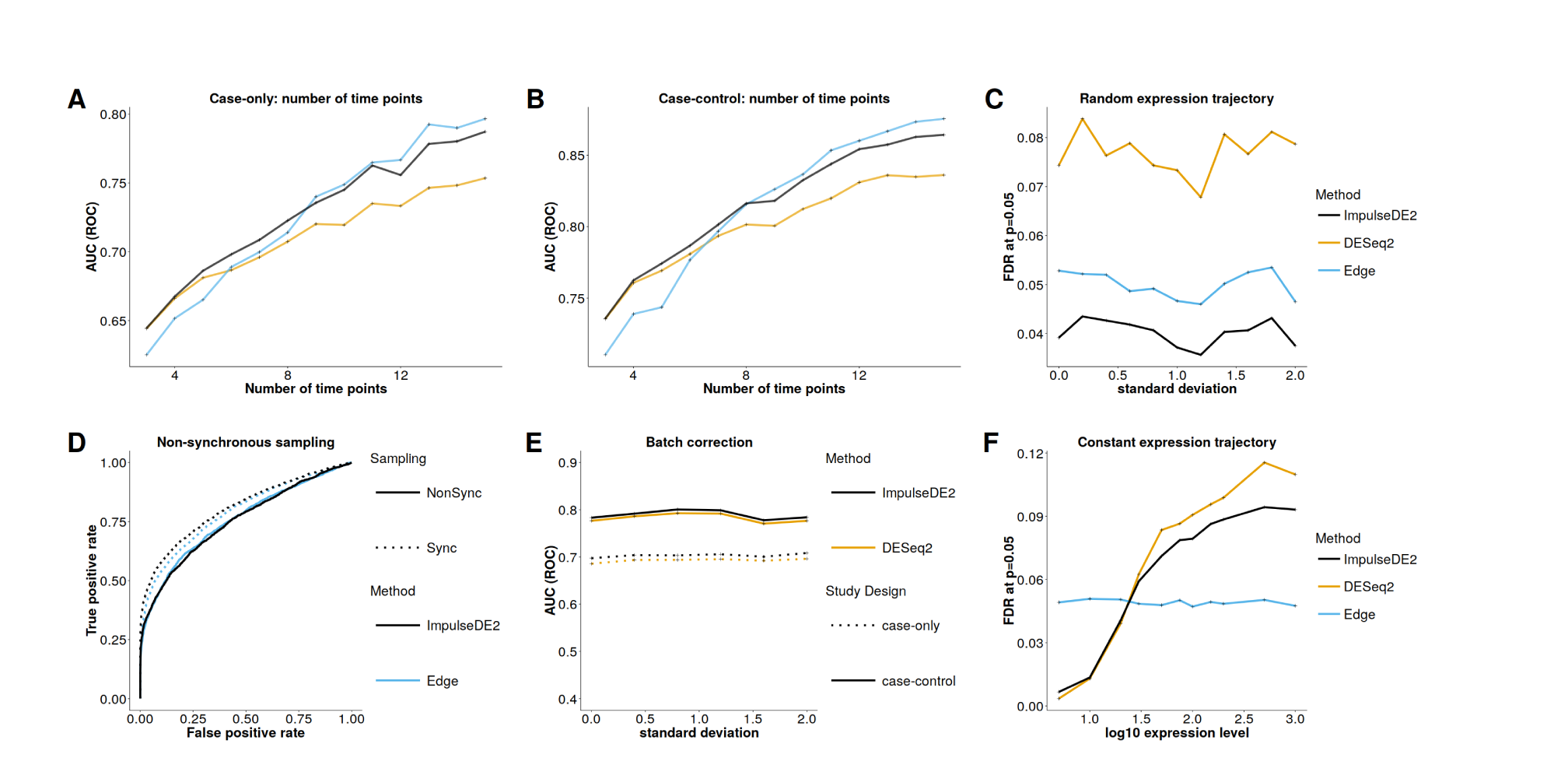
ImpulseDE2 unifies the individual advantages of DESeq2 and edge with respect to each other on simulated data. AUC: area under the curve, ROC: receiver-operator characteristic, FDR: false-discovery rate. Precision-recall curves for the presented data are in Fig. SI3. **A,B** ImpulseDE2 has higher statistical testing power than DESeq2 on data sets with many time points. AUC of ROC curve for case-only (**A**) and case-control (**B**) differential expression analysis with varying number of time points sampled. **C** ImpulseDE2 has a lower discovery rate on random expression trajectories than DESeq2. Sample means deviate from a constant expression trajectory by a factor drawn once for each time point from a normal distribution with mean one and the standard deviation indicated on the x-axis. The expression trajectory is therefore random with high variance if the the standard deviation of the normal distribution is high. We define every identified differentially expressed gene as a false discovery (Fig. 1D). **D** ImpulseDE2 is better suited for case-control analysis on “non-synchronously”-sampled conditions than DESeq2. ROC curve of ImpulseDE2 and edge for the case-control scenario with shared time points between samples of both conditions (Sync) and without shared time points (NonSync, “asynchronously”-sampled). The NonSync scenario cannot be easily handled with DESeq2. **E** ImpulseDE2 can correct for batch effects as well as DESeq2. AUC of ROC curve for case-only and case-control analysis with batch effects, the strength of the batch effects is quantified by the standard deviation of the normal distribution from which the batch factor is drawn. **F** ImpulseDE2 has a lower false discovery rate at very low expression levels than edge. All expression trajectories are constants with the expression value indicated on the x-axis. Every identified differentially expressed gene is a false discovery. Shown is the false discovery rate of each method for each range of simulated constant expression levels.

ImpulseDE2 and DESeq2 and less susceptible at high expression levels (Fig. 2F). Indeed, we found that edge called genes with low expression means differentially expressed which were not called differentially expressed by ImpulseDE2 in the data sets analyzed (Fig. 3E, 5C). This simulation suggests that these genes could be false-discoveries stemming from the noise model of edge.

**Figure 3:**
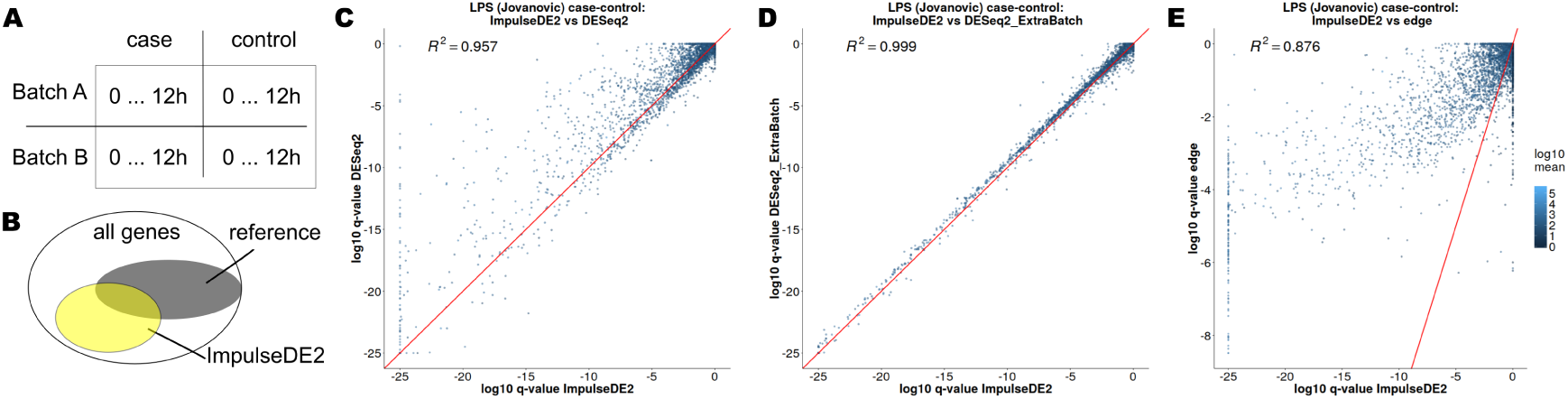
ImpulseDE2 out-performs edge and DESeq2 with a standard batch model on LPS (Jovanovic) case-control data. **A** Batch structure of data set: One replicate of each time point was obtained for each batch (A and B) and each condition (case and control). **B** Gene sets for gene enrichment analysis: Gene set of genes called differentially expressed by the reference method (grey) and by ImpulseDE2 (yellow). The set of genes only called differentially expressed by one method is the set of this method without the set intersection and was analyzed using gene set enrichment analysis. **C,D,E** Correlation plots of the inferred differential expression (case-control) Benjamini-Hochberg corrected p-values for all genes between ImpulseDE2 and DESeq2 with standard batch settings (**C**), DESeq2 with extra batch factor (**D**) and edge (**E**). *R*^2^ shown are Pearson correlations coefficients.

In summary, we show that ImpulseDE2 and edge have conceptual advantages over DESeq2 in studies with many time points and asynchronous sampling of case and control condition. Moreover, we show that ImpulseDE2 and DESeq2 are similarly good in batch correction and guarding against false-positives at low expression levels. Therefore, our simulations suggests that ImpulseDE2 combines the advantages of a continuous expression model with the advantages of the noise model of DESeq2.

### 1.3 ImpulseDE2 has advantages over DESeq2, edge and ImpulseDE on experimental data sets

We compare the performance of ImpulseDE2 with DESeq2, edge and ImpulseDE on four experimental data sets (LPS (Jovanovic), myeloid (Sykes), erythroid chromatin (Lara-Astiaso), hESC (Chu), as described in Fig. 1). We used ImpulseDE2 and DESeq2 on expected count matrices of the RNA-seq data sets. We used ImpulseDE and edge on size factor [2] (sequencing depth) normalized and log transformed expected count matrices. We present the results of the analysis of the LPS (Jovanovic) and myeloid (Sykes) data sets in the main text and the results of the analysis of hESC (Chu) and erythroid chromatin (Lara-Astiaso) data sets in the Supplementary Notes. All analysis performed is based on p-values and Benjamini-Hochberg corrected p-values. We did not use method specific false-discovery rate adjustment algorithms to make the results comparable.

#### 1.3.1 LPS (Jovanovic)

The LPS (Jovanvic) data set consists of duplicate samples of seven time points in two conditions (with and without LPS addition at time point 0h). The replicates come from two different batches (A and B) which cover both conditions (Fig. 3A). We observe batch effects between batches A and B in the correlation matrix of the samples (Fig. SI4).

We performed case-only and case-control differential expression analysis on the LPS (Jovanovic) data set. As expected, DESeq2 and ImpulseDE2 produce globally similar results on this data as the degrees of freedom used for the description of the mean trajectory are similar (Fig. 3, SI2). For case-control analysis, the standard batch model which would be used for DESeq2 is one batch correction factor per batch. ImpulseDE2 assign one batch correction factor per batch and condition as conditions are fit independently. We imitated this batch correction in DESeq2 (DESe2_ExtraBatch) (Supplementary Methods) and observed increased correlation between q-values between ImpulseDE2 and DESeq2 after this correction (Fig. 3C,D). The scatter plot of q-values of ImpulseDE2 and edge (Fig. 3 E) shows two groups of genes: Firstly, a group which receives higher q-values by ImpulseDE2 than edge which tends to have high expression means. Secondly, a group which receives low q-values by edge and is not called differentially expressed by ImpulseDE2 which tends to have very low expression means. We note that we also observed increased false-discovery rate of edge on genes with low expression means in simulations (Fig. 2F) suggesting that some of these genes could be false-positive differential expression calls by edge. We included plots of genes with strong differences in differential expression p-values between ImpulseDE2 and the reference methods in the supplement (Fig. SI20, SI21, SI22, SI23).

We performed gene set enrichment analysis with a hyper-geometric test against MSigDB [17] annotated gene sets to investigate the gene sets only labeled differentially expressed by ImpulseDE2 and not a reference method or only by the reference method and not by ImpulseDE2 (Fig. 3B, Supplementary Notes) at a q-value threshold of 1*e*— 2. Enrichment analysis associates a given gene set with annotated gene sets and can therefore be used to explore whether a set of potentially differentially expressed genes is related to the process in question. This is one way of approaching the unsolved problem of defining true positive differential expression labels in differential expression analysis on experimental data: A gene set is more likely to contain true positives if it is enriched in genes related to the underlying process.

Gene set enrichment analysis shows that the gene sets only called by ImpulseDE2 and not by the reference methods (either DESeq2 or edge) are enriched in immune signaling terms when compared with the MSigDB hallmark and GO biological process sets [18] (Supplementary Data 1,2). The sets only called differentially expressed by the respective reference method and not by ImpulseDE2 do not show as many and as strong enrichments related to the biological process (Supplementary Data 1,2).

We performed a similar comparison with ImpulseDE. ImpulseDE2 runs faster than ImpulseDE and shows better gene set enrichment results (Fig. SI5 and Supplementary Data 1).

To further validate the gene set enrichment analysis, we compared the number of overlaps of individual genes with gene sets from the ImmuneSigDB collection [19], which are related to dendritic cells, LPS or toll-like receptors. Each set in this collection corresponds to genes that are differentially expressed in a given comparison (e.g., stimulation with LPS vs. resting state). We interpret the number of overlaps as a ‘’responsiveness” metric for an individual gene: The more annotated gene sets related to the target process contain a gene, the more confident can we be in it being differentially expressed across the process. We analyzed the overall responsiveness of a set of genes only called differentially expressed by one method using the empirical density function of the number of overlaps. This analysis indicates that the sets only called differentially expressed by ImpulseDE2 are similarly relevant as the sets only called by edge or DESeq2 with the standard batch model, but are much larger (Fig. 4). We conclude, that ImpulseDE2 picks up more gene which are related to the underlying process than the edge and DESeq2.

**Figure 4:**
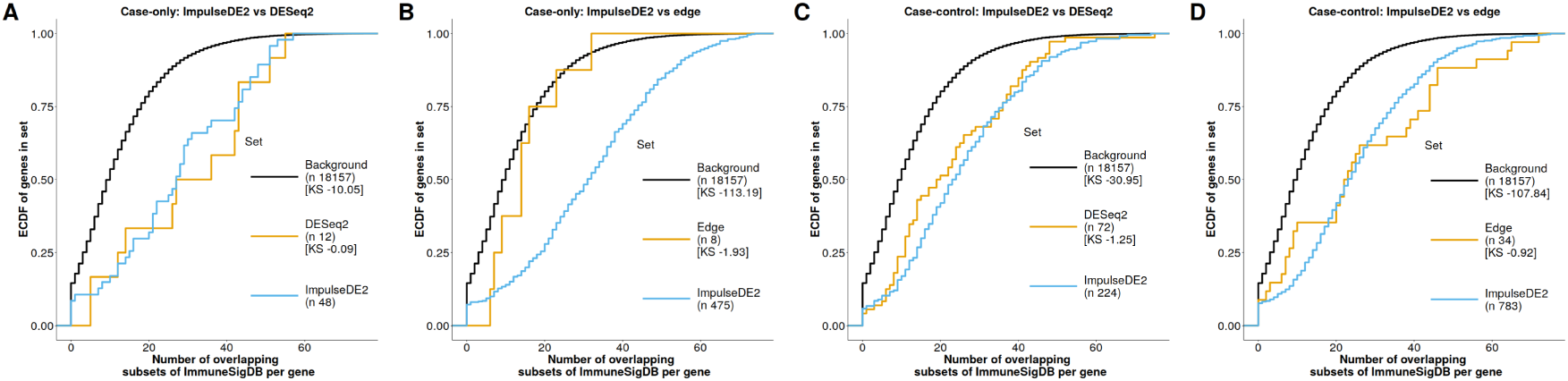
Overlaps to annotated gene sets indicate that ImpulseDE2 identifies more relevant genes than DESeq2 and edge on LPS (Jovanovic) data set. Background: All genes analyzed. ImpulseDE2: Genes only called differentially expressed by ImpulseDE2 and not by the reference method at an FDR-corrected p-value threshold of 0.01. Reference method (DESeq2 **A,C**, edge **B,D**): Genes only called differentially expressed by the reference method and not ImpulseDE2 at an FDR-corrected p-value threshold of 0.01. n: Gene set size. KS: log10 p-value of one sided Kolmogorov-Smirnov test for the empirical cumulative density functions (ECDF) of the “ImpulseDE2” set to lie below the ECDF of the “DESeq2” or “edge” set or below the “Background” set. The ECDF are based on the number of overlapping ImmuneSigDB [19] target sets with each gene in the individual gene sets: The target set was all ImmuneSigDB sets that contain any the following strings in their names: “DC”, “DENDRITIC” (empty, all listed under DC), “LPS” or “TLR”. **A,B** Case-only differential expression analysis. **D,E** Case-control differential expression analysis.

In summary, ImpulseDE2 is stable also on noisy data with batch effects and compares well against edge and ImpulseDE and as expected similar to DESeq2 as few time points were samples. Enrichment analysis shows that ImpulseDE2 is able to identify genes as differentially expressed which are meaningful for the underlying immune cell activation process and which are not identified by DESeq2, edge and ImpulseDE.

#### 1.3.2 Myeloid (Sykes)

Globally, ImpulseDE2 and DESeq2 give similar results on the myeloid (Sykes) data set (Fig. 5A,B). ImpulseDE2 has higher statistical testing power than DESeq2 on this data set because the number of time points (ten) is larger than the number of impulse model parameters (six). The advantage in testing power shows as a cloud of genes in the q-value scatter plot which lies above the diagonal (Fig. 5B). Moreover, ImpulseDE2 also catches 72 genes which contain observations with zero counts which are labeled as high variance outliers by DESeq2 and are therefore not regularized in the empirical Bayes dispersion estimation step of DESeq2. We note, that these genes tend to receive a variance that does visually not agree with the observations and are not called differentially expressed by by DESeq2 (Fig. 5B). Therefore, we implemented a correction step in ImpulseDE2 which automatically identifies genes with over-estimated variances and corrects the dispersion estimates to the maximum a-posteriori estimates from DESeq2. Because of the lower variance estimate, ImpulseDE2 can identify these genes as differentially expressed while they are missed by DESeq2 (Fig. 5B, Fig. SI6). However, there are also many genes which receive lower p-values by DESeq2 (bottom right half of Fig. 5B). We visually observed that these genes have mostly non uni-modal trajectories (Fig. SI7). Non-constant and non uni-modal trajectories could be multi-modal or oscillating, such multi-modal patterns are however difficult to resolve with such a sampling frequency. The potentially multi-modal patterns are indeed visually difficult to distinguish from randomly generated mean signals for which we previously established on simulated data that ImpulseDE2 penalizes these as noise while DESeq2 labels these trajectories as differentially expressed (Fig. 2C).

**Figure 5:**
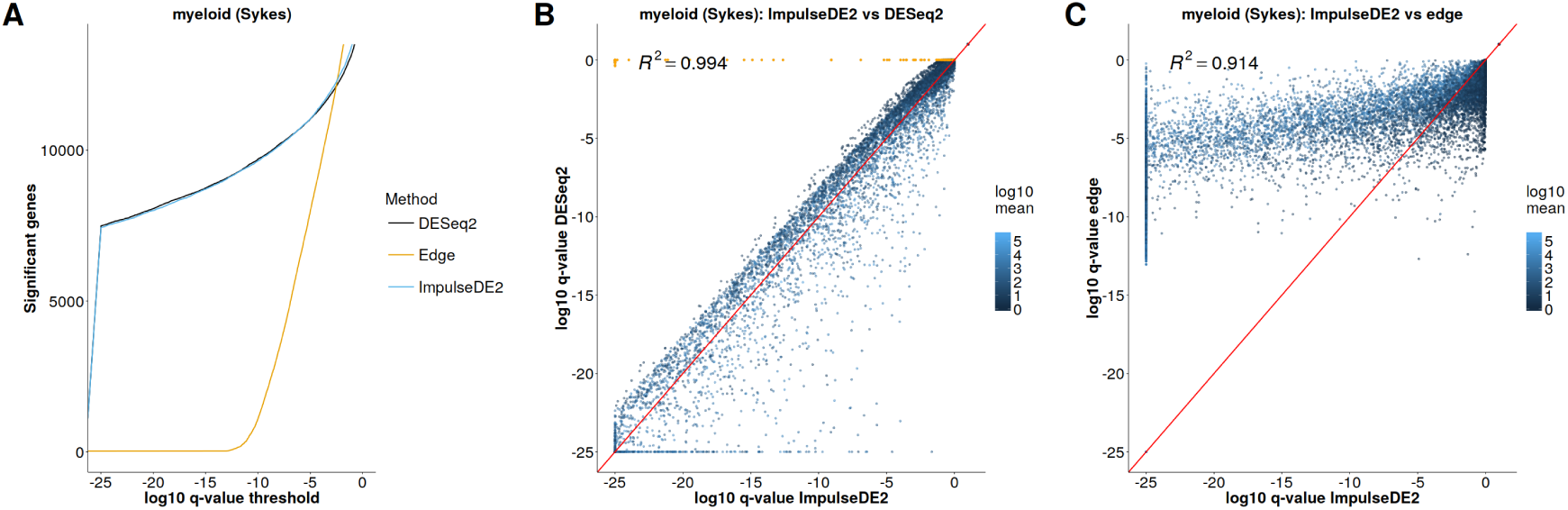
Figure 5: ImpulseDE2 out-performs DESeq2 and edge on myeloid (Sykes) data set. Orange points correspond to genes for which ImpulseDE2 used the DESeq2 maximum a-posteriori dispersion estimate and disabled DESeq2 dispersion outlier handling. **A** Number of significantly differentially expressed genes as a function of the significance threshold.**B** Correlation plot of the inferred differential expression Benjamini-Hochberg corrected p-values for all genes between ImpulseDE2 and DESeq2. **C** Correlation plot of the inferred differential expression Benjamini-Hochberg corrected p-values for all genes between ImpulseDE2 and edge. R^2^ shown are Pearson correlations coefficients.

The gene set only called differentially expressed by ImpulseDE2 and not by DESeq2 is over-represented in sets associated with immune response and immune signaling when compared with the GO biological process sets [18] (Supplementary Data 4). The set only called differentially expressed by DESeq2 and not by ImpulseDE2 is over-represented in membrane vesicle terms. Myeloid differentiation is immune cell development which suggests the immune related terms are more likely true positive calls than membrane vesicle terms. We interpret the enrichment of the set of genes only called differentially expressed by ImpulseDE2 and not by DESeq2 in immune related GO terms as an enrichment of true positive differential expression calls in this gene set: ImpulseDE2 finds truely differentially expressed genes which are missed by DESeq2. At the same time, ImpulseDE2 is more robust than DESeq2 as the genes missed by ImpulseDE2 do not have a clear trajectory structure and are not enriched in relevant biological terms.

We observed large differences in the global differential expression results of ImpulseDE2 and edge on the myeloid (Sykes) data set (Fig. 5A,C). Many of the genes only labeled differentially expressed by edge have low mean expression (Fig. 5C, SI9) and visually appear much less convincing than the genes only labeled differentially expressed by ImpulseDE2 (Fig. SI8, SI9) which suggests that these genes only called differentially expressed by edge could be false-positives stemming from the edge noise model (Fig. 2F). We did not see striking enrichment results in the comparison between ImpulseDE2 and edge (Supplementary Data 4).

In summary, ImpulseDE2 has advantages over DESeq2 due to higher statistical testing power, robustness to fluctuations and variance outlier detection and over edge due to its noise model on the myeloid (Sykes) data set. Similarly, we found that ImpulseDE2 outperforms ImpulseDE on this data set (Fig. SI5B, Supplementary Data 4).

### 1.4 ImpulseDE2 identifies genes with transiently changing expression level

The response of a cell to a stimulus can often be viewed as a transition of the population from one transcriptomic state to another transcriptomic state, such as in cell activation or differentiation [9]. A biologically more interesting question than simple differential expression may be, whether a gene is responsible for the different phenotypes of the transcriptomic states or whether it drives the transition between the states. We introduce a hypothesis testing scheme that is able to answer these questions. ImpulseDE2 can fit a sigmoid model in addition to the impulse and constant model to each gene. This sigmoid models a monotonous expression trajectory. We define transiently regulated genes as genes which are significantly better fit by an impulse model than by a sigmoid model and which do not have a monotonous impulse model fit. We define permanently regulated genes as genes which are not transient but which are significantly better fit by a sigmoid than by a constant model. Supplementary Data We classify up- and down-regulated genes in the transient and the permanent class based on their impulse model fits. This simple approach does not consider the permanent effect which transiently up- or down-regulated genes may have: Different initial and final states. One may easily formulate alternative tests based on a constant, a sigmoid and an impulse model fit which consider these effects.

#### 1.4.1 hESC (Chu)

The heatmap of expression profiles sorted by their peak times within each class shows that ImpulseDE2 can indeed classify expression trajectories into transiently and permanently changing trajectories (Fig. 6). The permanently down-regulated genes indicate that the embryonic state is left between 0 and 12 hours. The permanently up-regulated genes indicate that final state is reached around 72 hours. Both times are consistent with the PCA of single-cell RNA-seq samples of the same system [14].

**Figure 6:**
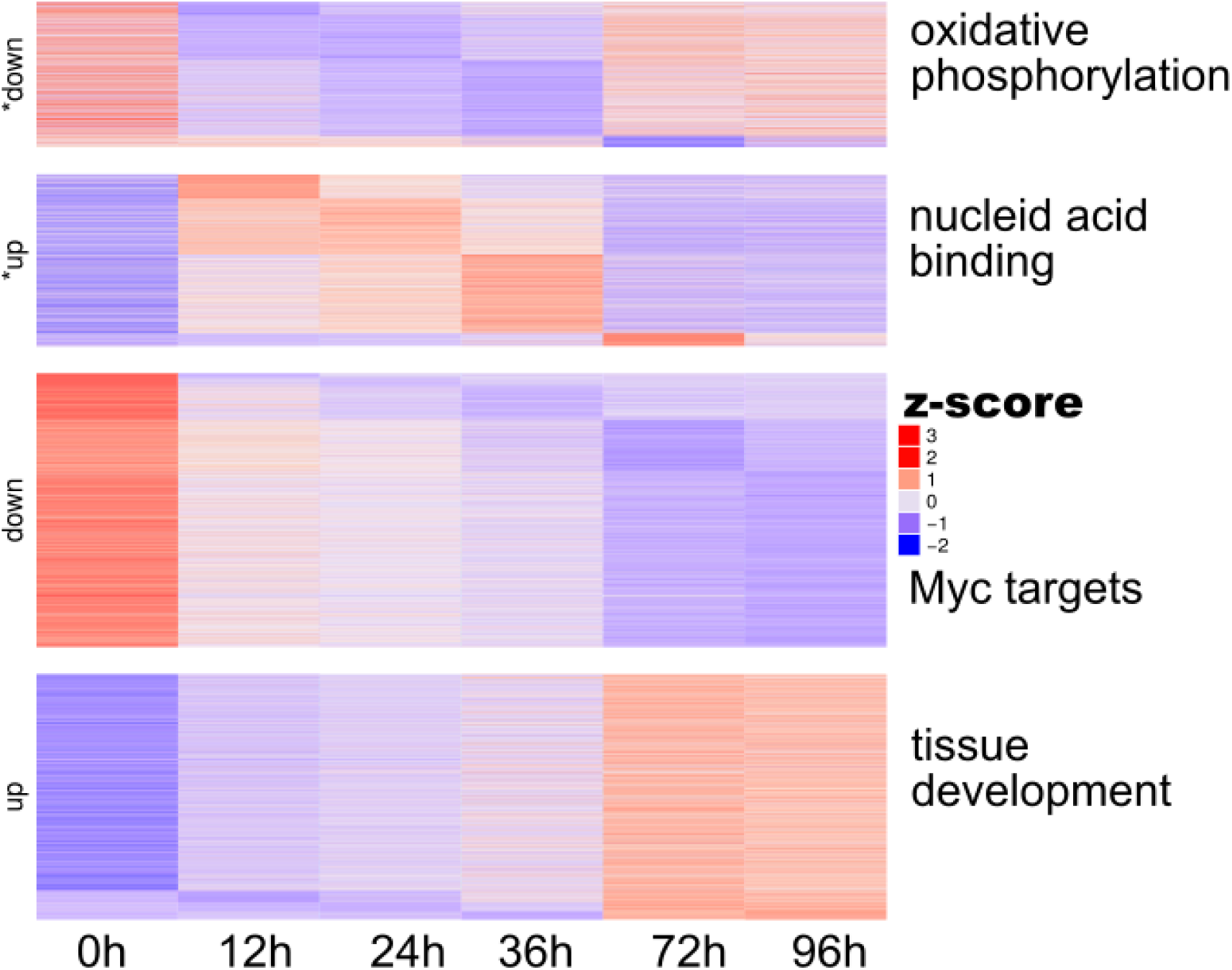
ImpulseDE2 can distinguish between transiently and permanently changing expression trajectories in hESC (Chu) data set. *up: transiently increased trajectories. *down: transiently decreased trajectories.up: monotonously (permanently) increasing trajectories. down: monotonously (permanently) decreasing trajectories. Shown are z-scores of size-factor normalized mean expression values per time point. Selected top enrichments with GO biological process, GO molecular function and MSigDB hallmark gene sets are supplied to the right of each group.

We find a large group of genes which is transiently down regulated from 0 to 72 hours after differentiation induction. The top gene set enrichment hit of the transiently down regulated genes against the GO molecular function gene sets [18] is oxidoreductase_activity and the top enrichment with respect to the C2 hallmark set is oxidative phosphorylation (Supplementary Data 5). Chu et al. also show that the human embryonic stem cells can be induced to differentiate by hypoxic conditions but did not use hypoxic condition in the presented data set hESC (Chu). We argue that the observed down-regulation of genes related to oxidative phosphorylation during differentiation may provide a mechanistic link to the differentiation induction by hypoxia: Down-regulation of oxidative phosphorylation related processes is a molecular (for example transcriptomic or metabolic) signature that is induced by and drives differentiation. Therefore, differentiation can be directly induced by inducing this signature through hypoxia. We note that the optimal hypoxia treatment for differentiation induction coincides with the time frame of down regulation of the oxidative phosphorylation related gene set (0 to 72h, [14] and Fig. 6).

Moreover, we find a succession of transiently up-regulated genes as would be expected as result of a signaling cascade that drives the transition from initial state to final state (reached at 72 hours). Indeed, the top gene set enrichment hits of these transiently up-regulated genes against the GO molecular function gene sets contain many nucleic acid binding terms, suggesting gene expression regulation cascades are active.

The top three enrichments of permanently down-regulated genes against the MSigDB hallmark gene sets [20] contain two Myc-target sets which suggests, that this gene set represents the loss of embryonic cell identity. The top enrichments of permanently up-regulated genes against the GO biological process gene sets [18] contain several tissue development terms which suggests that this gene set represents the gain in differentiated cell identity.

In summary, we find that genes with transient expression trajectories reflect transient processes in the population and genes with monotonous expression trajectories to reflect differences in the cell states (embryonic and definite endoderm).

## 2 Discussion

We motivated the use of the impulse model by showing that the impulse model is suitable to model transcriptomic and epigenomic dynamics of cells in response to environmental and developmental stimuli.

The impulse model has previously been used in the differential expression tool ImpulseDE which operates on a model-free noise distribution. Here we introduce ImpulseDE2, a differential expression algorithm which is based on the impulse model and tailored to count data. The altered noise model makes ImpulseDE2 faster than ImpulseDE and yields better differential expression results on count data. Moreover, ImpulseDE2 has conceptual advantages over DESeq2 on longitudinal studies if many time points are sampled or time points are sampled asynchronously between case and control condition. We show on simulated and experimental data sets that ImpulseDE2 penalizes multi-modal expression trajectories relative to DESeq2, resulting in lower p-values for differential expression assigned by ImpulseDE2 compared to DESeq2. We argue that such multi-modal trajectories are difficult to distinguish from a noisy background level and analysis may benefit from omitting these trajectories. Gene set enrichment analysis supports the differential expression calls of ImpulseDE2. ImpulseDE2 gives consistently better results than edge and ImpulseDE on count data.

ImpulseDE2 relies on estimation of a non-linear model for expression as a function of time. Disadvantages of such models include numerical estimation problems and local maxima of the log-likelihood cost function. We guarded against both problems in the implementation through multiple initializations and through numerical thresholds. ImpulseDE2 did not produce numerical errors on any analyzed data set. We did not observe model fits which do not capture the observed uni-modal patterns in any of the discussed data sets.

One major advantage of ImpulseDE2 is that it has high statistical testing power on data sets with many time points. Many “time points” are sampled in pseudotemporal projections of single-cell RNA-seq data sets, where time is replaced by pseudotime which is a measure for developmental progression. Therefore, continuous models that can faithfully represent the global expression dynamics in such systems are needed. We propose the impulse model as one suitable model for such systems.

Finally, we combine impulse model fits with constant and sigmoid model fit to identify genes with transiently or monotonously changing trajectories and show that these automatically annotated gene sets represent biologically meaningful groups of genes. Our analysis suggests a mechanism for hypoxia-induced human embryonic stem cell differentiation.

## 3 Methods

ImpulseDE2 fits an impulse [21] model to time course count data and performs differential expression analysis based on the model fits with a log-likelihood ratio test. The central co-variate considered in ImpulseDE2 is continuous time. Secondly, if trajectories from two different conditions (case and control) are compared, a discrete condition indicator co-variate is added. Thirdly, if batch structure is present in the data beyond the case and control conditions, a categorial batch assignment co-variate is added for each confounding variable.

We distinguish case-only and case-control differential expression (DE) analysis. Case-only DE analysis is DE analysis over time. In case-only DE analysis, an impulse fit to the expression values over time is compared against a constant fit. The goal of case-only DE analysis is to find genes which behave nonconstant over time. Case-control DE analysis is DE analysis between time trajectories of samples from two conditions, such as a case and control condition in a treatment study. In case-control DE analysis, separate impulse model fits to each condition are compared against an impulse model fit to all samples combined. The goal of case-control DE analysis is to find genes which have differential temporal behavior between the two conditions.

ImpulseDE2 can additionally correct for batch effects through a gene and batch specific factor in the gene expression model. Multiple confounding variables with differing batch structures can be modeled if the corresponding design matrix is full rank.

### 3.1 The impulse model

ImpulseDE2 models the expression level of a gene as a function of time with the impulse model *f*_Impulse_. The impulse model is the scaled product of two sigmoid functions (eq. 1) [21] and has three states with a state-specific expression value each: initial, peak and steady state. The two sigmoid functions represent the transitions of initial state to peak state and peak state to steady state.

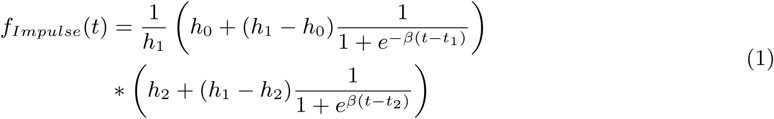

*h*_0_, *h*_1_ and *h*_2_ are the initial, peak and steady state expression values (amplitude parameters). *t*_1_ and *t*_2_ are the state transition times, *β* is the slope parameter of both sigmoid functions. One could use two different slope parameters but we use a shared slope parameter to minimize degrees of freedom of the alternative model in case-only differential expression analysis.

### 3.2 The sigmoid model

ImpulseDE2 can model the expression level of a gene as a function of time with the sigmoid model *f_sigmoid_.* The sigmoid model is nested within the impulse model and represents a simplified monotonous state transition.

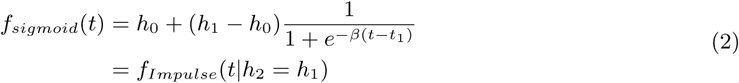

*h*_0_ and *h*_1_ are the initial and steady state expression values (amplitude parameters). *t_1_* ais the state transition times, *β* is the slope parameter of the sigmoid function.

### 3.3 The likelihood function

We assume that the number of reads x generated from *μ* transcripts is negative binomially distributed. The likelihood (𝓛x_i,._|*μ*_i,._,*ϕ*_i_) of the count data *x_i_*,. of gene *i* observed in *J* samples at time points *t(j)* is:

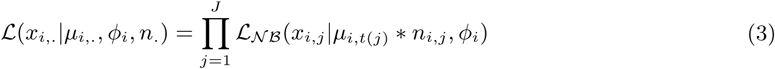

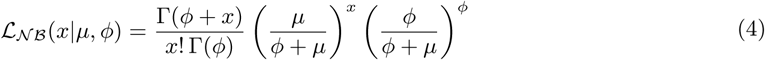

where*𝓛_𝒩ℬ_* is the negative binomial likelihood. Per gene, the proposed negative binomial model has one mean parameter *f*μ*_i,t(j)_* per time point observed in a condition, one normalization factor *n_i_*,_*j*_ per gene and sample and one dispersion factor ϕ_i_ for all samples of a gene. The negative binomial dispersion parameter links the mean of a negative binomial distribution to its variance: Overdispersion relative to the Poisson process is modeled with positive dispersion factors.

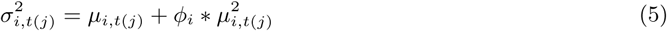

The normalization factor scales the model prediction for an observation according to a sample-specific size factor *s_j_* and a gene- and sample-specific batch factor *b_i_,_l(j)_* for each confounding variable *l*:

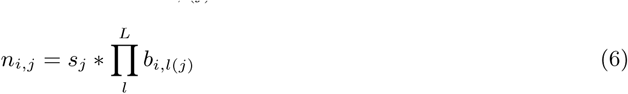

where*l*(*j*) is the batch index of sample *j* with respect to the confounding variable *l* out of *L* confounding variables. The size factor reflects differences in sequencing depth between samples. The batch factor reflects relative differences in the amount of transcripts in corresponding samples of different batches and is defined for each gene. Different confounding variables could for example be patient identity and sequencing batch (refer to section Supplementary Methods 2.1 for an example). The design matrix of the confounding factors has to be full rank: Any confounder may not be a linear combination of the other confounders.

The overall likelihood of the data is the product of the gene-wise likelihoods. Therefore the optimization of the model parameters based on the likelihood intuitively lends itself to parallelization over genes.

### 3.4 ImpulseDE2 algorithm

Input Matrix of count data (*N* genes x *J* samples) and table specifying meta data of samples.

1. **Hyper-parameter estimation:** Run DESeq2 and save dispersion parameters. Catch high variance dispersion parameter outliers. Compute size factors.
2. **Model fitting:** If performing case-control differential expression analysis: Split data set into three data sets with only samples contained in case condition (”case”), only samples contained in control condition (”control”) and with all samples (”combined”) and iterate over data sets. Parallelize over genes within a data set.
  a. **Constant model:** Optimize constant model parameters based negative binomial likelihood with the Broyden–Fletcher–Goldfarb–Shanno algorithm (BFGS) (up tp I iterations).
  b. **Sigmoid model:** Optional, only fit if transients trajectories are to be identified.
    i. **Initialization:** Initialize two sigmoid models for each gene based on a increase and a decrease model.
    ii. **Optimization:** Optimize sigmoid model parameters of both initializations based on negative binomial likelihood with BFGS with the dispersion parameter estimated by DESeq2 (up tp I iterations).
    iii. **Fit selection:** Keep sigmoid model fit with highest log-likelihood.
  c. **Impulse model:**
    i. **Initialization:** Initialize two impulse models for each gene based on a peak and a valley model (explained below in Model fitting, Fig. 1A).
    ii. **Optimization:** Optimize impulse model parameters of both initializations based on negative binomial likelihood with BFGS with the dispersion parameter estimated by DESeq2 (up tp I iterations).
    iii. **Fit selection:** Keep impulse model fit with highest log-likelihood.
3. **Differential expression analysis:** Perform loglikelihood ration test. Compute p-values from χ^2^- distributed deviance and perform false-discovery rate correction.

**Output** Benjamini-Hochberg false-discovery rate corrected p-values for differential expression for each gene. If selected, a classification of differentially expressed genes into transiently and monotonously changing expression trajectories.

The algorithmic complexity the run time dominating impulse fitting step of ImpulseDE2 is *𝒪(I* * *N* \**J*). The run time of ImpulseDE2 is approximately linear in the number of genes and samples. Therefore, we expect ImpulseDE2 to scale well to large data sets. The number of numerical fitting iterations necessary to reach convergence may however differ between data sets and increase with the number of samples leading to above linear complexity in the number of samples. ImpulseDE2 is parallelized over genes in the run-time dominating model fitting steps (constant, impulse and sigmoid model).

### 3.5 Hyper-parameter estimation

Hyper-parameter estimation precedes the mean model fitting step in ImpulseDE2. During hyper-parameter estimation, sample-specific normalization factors (size factors) and gene-specific dispersion parameters are estimated. The estimation of the mean model (impulse, sigmoid and constant) is conditioned on the hyperparameters.

#### 3.5.1 Dispersion parameters

Dispersion parameters have been previously regularized by smoothing gene-wise point estimators of genes according to a genome-wide trend in an empirical Bayes scheme [1]. Such a smoothing breaks the independence of gene-wise expression models. To maintain independence between the gene-wise expression models, we estimate the dispersion parameters as hyper-parameters prior to fitting the mean model. The dispersion parameters are estimated with DESeq2 [2] treating time as a categorial variable. ImpulseDE2 catches high-dispersion outliers in genes which contain zero count observations which are not regularized through empirical Bayes smoothing by DESeq2 and replaces the raw dispersion estimate with the a-posteriori estimate from DESeq2. This outlier handling results in much lower regularized variance estimates of the outlier genes and does not affect the remaining genes.

#### 3.5.2 Size factors

The normalization factor *s_j_* of sample *j* is the median of the ratios of observed counts within this sample to the geometric means *κ_i_* of the genes *i* [2]. Size factors are computed on the subset of genes which do not contain zero observations *I_\0_.*

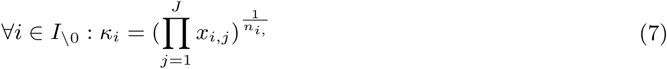

where*n_i}_.* is the number of observations for gene *i.*

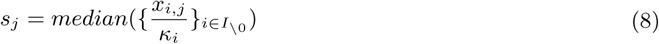

The median ratio of observed counts to the geometric mean over all genes *I* has been previously used as a sample normalization factor because it is less sensitive to regions with high mean and high variance count distributions than the sequencing depth [2].

### 3.6 Model Fitting

#### 3.6.1 Constant model

The constant mean parameter is estimated as a maximum likelihood estimator based on a negative binomial loglikelihood with the dispersion factor as the hyper-parameter inferred using DESeq2. The optimization is performed with the BFGS algorithm.

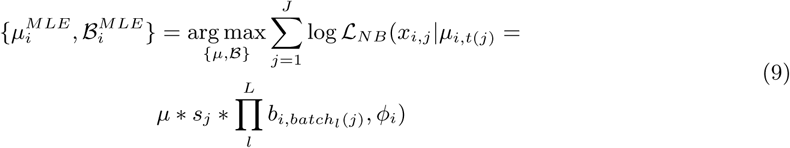

where *μ*_*i*_ is the constant mean parameter of gene *i* and *B_i_*={{b_*i*_,ι_κ}*κ*∈*1:K*}_ iιєgi:L is the set of batch factors for gene i. Note that one vector of batch factors is estimated for each confounding variable. *b_i_l_1_*ι_*l*_ is one because *k*= 1 is the reference batch in each condition and for each confounder.

#### 3.6.2 Impulse model

##### Initialization

Each gene model is initialized twice based on a peak and a valley model. Firstly, one expression level is estimated for each time point by averaging all size factor-corrected samples. If a batch structure is given, the expression means are corrected for batch effects based on batch factor estimates. The batch factors are estimated as the ratio of the mean size factor-corrected expression level of the samples of a given batch of a given gene and all samples of this gene. Initial (h0) and steady state (h2) are initialized as the expression level at the first and the last time point. The transition state is estimated as the highest (peak) or lowest (valley) expression level between first and last time point (’extremum’). To estimate the transition time parameters t1 and t2, expression gradients are locally linearly approximated between adjacent time points. t1 is initialized as the average of the two time coordinates associated with the maximal (peak) or minimal (valley) linear gradient approximation out of the time points before the estimated ‘extremum’. t2 is initialized as the average of the two time coordinates associated with the minimal (peak) or maximal (valley) linear gradient approximation out of the time points after the estimated ‘extremum’.

##### Optimisation

The cost function for the fit is the negative binomial loglikelihood of the data, the value of the impulse model is the mean parameter. The optimization is performed with the BFGS algorithm.

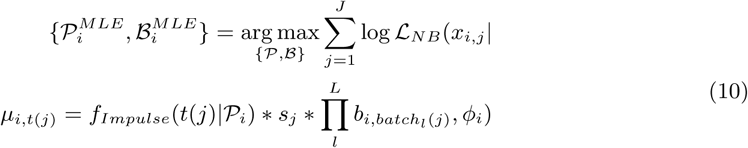

where *𝒫_i_*={*h*_o_,*h*_1_, *h*_2_, *t*_1_, *t*_2_,*β*} is the set of parameters of the impulse model fit to gene i and *ℬ_i_*={*{b_i_,l_κ_}κ∈ 1:K}l*∈_*1:L*_ is the set of batch factors for gene *i*. *b_i,1_* is one because *k*= 1 is the reference batch in each condition. *f_Impuls_s(t|𝒫_i_)* is the impulse model with the parameters *𝒫_i_* evaluated at time point t. ImpulseDE2 fits the amplitude parameters of the impulse model and batch factors in log space and imposes a loglikelihood sensitivity boundary at 10^−10^ below which changes in the parameters do not affect the loglikelihood which constraints the value of the impulse function to be positive. Thereby, the negative binomial mean parameter is guaranteed to be larger than zero.

##### Sigmoid model

Fit as the impulse model but with the sigmoid instead of the impulse model (eq. 2). Initializations are based on a monotonous increase and a monotonous decrease model

### 3.7 Differential expression analysis

#### 3.7.1 Standard differential expression analysis

Model selection between alternative and null model is based on a loglikelihood ratio test. P-values for differential expression are computed based on the χ^2^-distributed deviance and are false-discovery rate corrected [22]. Note that the deviance is only χ^2^-distributed if the null model is nested within the alternative model which is given in both case-only and case-control differential expression analysis (proof in Supplementary Notes).

#### 3.7.2 Identification of transiently activated genes

Again, model selection between alternative and null model is based on a loglikelihood ratio test. P-values for differential expression are computed based on the χ^2^ distributed deviance and are false-discovery rate (FDR) corrected [22]. Transiently regulated genes are defined as genes which have FDR-corrected p-value of the comparison impulse model against sigmoid model below a significance threshold. Moreover, transiently regulated genes are additionally required to not have a monotonous impulse model fit. Genes with monotonous expression trajectories are defined as genes which do not meet one of the above two conditions but which have a FDR corrected p-value of the the comparison sigmoid model against constant model below a significance threshold. Up- and down- regulation is defined based on the impulse model fits (Fig. 1A): Up-regulated transient genes have a peak-like trajectory, down-regulated transient genes have a valley like trajectory, permanently up-regulated genes have a monotonously increasing trajectory, permanently down-regulated genes have a monotonously decreasing trajectory.

